# Long reads and Hi-C sequencing illuminate the two compartment genome of the model arbuscular mycorrhizal symbiont *Rhizophagus irregularis*

**DOI:** 10.1101/2021.08.12.456011

**Authors:** Gokalp Yildirir, Jana Sperschneider, Malar C Mathu, Eric CH Chen, Wataru Iwasaki, Calvin Cornell, Nicolas Corradi

**Affiliations:** Department of Biology, University of Ottawa, ON, Ottawa, K1N 6N5, Canada; Biological Data Science Institute, The Australian National University, Canberra, Australia; Department of Integrated Biosciences, Graduate School of Frontier Sciences, The University of Tokyo, Japan

## Abstract

Chromosome folding links genome structure with gene function by generating distinct nuclear compartments and topologically associating domains (TADs). In mammals, these undergo preferential interactions and regulate gene expression. However, their role in fungal genome biology is unclear. Here, we combine Nanopore (ONT) sequencing with chromatin conformation capture sequencing (Hi-C) to reveal chromosome and epigenetic diversity in a group of obligate plant symbionts; the arbuscular mycorrhizal fungi (AMF). We find that five phylogenetically distinct strains of the model AMF *Rhizophagus irregularis* carry 33 chromosomes with substantial within species variability in size, as well as in gene and repeat content. Strain-specific Hi-C contact maps all reveal a ‘checkerboard’ pattern that underline two dominant euchromatin (A) and heterochromatin (B) compartments. Each compartment differs in the level of gene transcription, regulation of candidate effectors and methylation frequencies. The A-compartment is more gene-dense and contains most core genes, while the B-compartment is more repeat-rich and has higher rates of chromosomal rearrangement. While the B-compartment is transcriptionally repressed, it has significantly more secreted proteins and *in planta* up-regulated candidate effectors, suggesting a possible host-induced change in chromosome conformation. Overall, this study provides a fine-scale view into the genome biology and evolution of prominent plant symbionts, and opens avenues to study the epigenetic mechanisms that modify chromosome folding during host-microbe interactions.

## Introduction

Arbuscular mycorrhizal fungi (AMF) are obligate plant mutualistic organisms of the fungal subphylum Glomeromycotina (Spatafora *et al*., 2016) that evolved a symbiotic relationship with the roots of most vascular plants species (Corradi & Bonfante, 2012; Martin *et al*., 2017; Lutzoni *et al*., 2018). During the AM symbiosis, these fungi enhance their host nutrient and water uptake in exchange of sugars and lipids, and provide an increased resistance against pathogens (Bonfante & Anca, 2009). The first fossil records of the AM symbiosis date several hundred million years, and today this symbiotic relationship between AMF and their plant hosts is widespread globally, highlighting AMF’s capacity to persist in various environmental conditions (Delaux & Schornack, 2021). AMF spores and hyphae harbor up to thousands of coexisting nuclei at all times (Kokkoris *et al*., 2020), and their genetics are defined by the presence of two life stages – i.e. the homokaryotic stage (AMF homokaryons), where co-existing nuclei are genetically largely uniform, and the heterokaryotic stage (AMF dikaryons), where nuclei originating from two parental strains coexist in the mycelium (Ropars *et al*., 2016; Corradi & Brachmann, 2017; Chen *et al*., 2018a; Kokkoris *et al*., 2021).

Genome analyses revealed that species in the Glomeromycotina carry gene losses in cellular pathways that inhibit the metabolism of fatty acids, sugars and plant cell wall degradation enzymes (Tisserant et al., 2013; Lin et al., 2014; Chen et al., 2018b; Morin et al., 2019; Malar C et al., 2021a), and contain an abundance of transposable elements (TE). Closely related AMF strains also vary dramatically in their gene content, suggesting that AMF have pangenomes (Chen *et al*., 2018b; Mathieu *et al*., 2018a). The pangenome concept highlights the distinction between ‘core’ conserved genes hypothesized to be essential for daily activities (e.g. cytoskeleton, protein translation, etc.), and ‘dispensable’ non-conserved genes thought to help individual strains adapt to changing environments and hosts (e.g. transduction signalling, effectors, etc.) (Li *et al*., 2018; McCarthy & Fitzpatrick, 2019; Badet *et al*., 2020) (Li *et al*., 2018; McCarthy & Fitzpatrick, 2019; Badet *et al*., 2020).

Identifying the mechanisms from which pangenomes emerge is essential for understanding the mode of evolution of these widespread plant symbionts. In fungal plant pathogens, high repeat content generates within-species genome variability and it was proposed that similar mechanisms drive genetic variability in AMF (Chen *et al*., 2018b; Mathieu *et al*., 2018a). However, this hypothesis is currently untestable for AMF because most available genome assemblies are highly fragmented – i.e. composed in many cases of dozens of thousands of contigs (Chen *et al*., 2018b; Mathieu *et al*., 2018b). This high fragmentation rate can generate mosaic sequences of multiple haplotypes (collapsed regions), chimeric assemblies and artificial gene duplications (Denton *et al*., 2014; Montoliu-Nerin *et al*., 2020), preventing fine-scale and reliable analyses of genome rearrangements, repeat diversity, and repeat plasticity within and across strains.

The highly fragmented state of AMF genomes also hampered the identification of mechanisms that define relationships between genome structure and function. Within nuclei, chromosomes separate into two groups: active open chromatin A-compartments (euchromatin) and less active chromatin B-compartments (heterochromatin) and each compartment can be differently regulated (Pombo & Dillon, 2015; Dekker & Heard, 2015; Szabo *et al*., 2019; Kempfer & Pombo, 2020). This compartment-driven gene regulation highlights the key epigenetic role of chromosome folding for genome biology, but knowledge of this process in fungi was thus far restricted to a few model genera (e.g. *Sacharomyces, Neurospora*) and the filamentous fungus *Epichloë festucae* (Galazka *et al*., 2016; Kim *et al*., 2017; Winter *et al*., 2018).

In the plant symbiont *E. festucae*, chromosome folding and repetitive elements together divide the genome into distinct regions with very similar gene expression suggesting that both genome characteristics co-regulate symbiotic associations in this species. However, whether similar mechanisms drive gene expression and symbiotic associations in other fungal symbionts, including AMF, is unknown. Here, we aimed to obtain a complete understanding of the AMF genome biology and within species genome dynamics by acquiring long-read and strain-specific high-throughput chromatin conformation capture (Hi-C) sequencing data (Schmitt *et al*., 2016) from five phylogenetically distinct homokaryotic strains of the species *R. irregularis*.

## Materials and Methods

### Culturing, DNA Extraction and ONT/Hi-C Sequencing

*R. irregularis* strains A1 (DAOM-664342), B3 (DAOM-664345) and C2 (DAOM-664346) were originally isolated from Switzerland (Tänikon), while the strain DAOM-197198 was originally isolated from Pont Rouge, Canada (Koch *et al*., 2004; Stockinger *et al*., 2009) and the strain 4401 (DAOM-240446) was isolated from *Ammophila breviligulata* in “La Martinique Îles-de-la-Madeleine” (Canada).

All five strains were cultured *in vivo*, using *Daucus carota* root organ cultures (ROCs) as the host, as previously described (Corradi *et al*., 2004). The strains propagated in two-compartment ROCs, allowing us to produce mycelium and spores without obvious contaminants. For Oxford Nanopore (ONT) sequencing, high quality, high molecular weight DNA was extracted for the strains A1, C2, B3 and 4401 using a protocol proposed by Schwessinger (Schwessinger, 2016). Following extraction, high quality DNA samples were processed using Nanopore Ligation Sequencing Kit SQK-LSK109 to prepare the sequencing libraries, which were sequenced using the MinION R9.4.1 flowcells to produce an average of 8.45 million reads with an N50 > 4Kb, reaching to a genome coverage of 160x.

### Chromosome assembly and annotation

ONT reads were basecalled with guppy (version 4.5.2) using the “high accuracy” configuration (dna_r9.4.1_450bps_hac.cfg), and were used to generate assemblies using Canu (version 2.0) (Koren *et al*., 2017). These assemblies were then further polished using Racon (v 1.4.10) (Vaser *et al*., 2017) using the default parameters and the consensus sequences were created using Medaka (version 1.0.3). The polishing step was concluded with two rounds of Pilon (version 1.23) using Illumina reads and default parameters (Walker *et al*., 2014). Illumina reads were then mapped on the polished contigs using bwa mem (version 0.7.17) (Li, 2013) and the read coverage of each contig was calculated using bedtools (version 2.26.0) (Quinlan & Hall, 2010). Contigs having abnormally low (<5x) and high (>200x) coverages were filtered out using purge_haplotigs (Roach *et al*., 2018), and the remaining contigs were used for further scaffolding. Illumina reads used for polishing from the strains A1, B3 and C2 were obtained from Ropars *et al*. (Ropars *et al*., 2016), DAOM197198 reads were obtained from Chen et al. 2018(Chen *et al*., 2018b), 4401 Illumina reads were used in this study for the first time.

Approximately 200 mg of fresh mycelium from each strain was crosslinked and shipped to Phase Genomics (Seattle, WA) to obtain strain-specific Hi-C data. Hi-C data was produced using the Proximo Hi-C Kit (Microbe) at the Phase Genomics with DPNII restriction enzymes. Before generating high-throughput Hi-C Illumina data, the quality of these libraries was assessed by mapping a low coverage paired-end data onto available *R. irregularis* assemblies. A library is deemed to be of high quality if the fraction of high quality paired-end reads mapping respectively > 10KB apart within and across publicly available contigs exceeds 1.5% and 5.0%. In all cases, the values obtained for our samples far exceeded the QC limits set by Phase Genomics (Seattle, WA) – e.g. average 11.75% and 23%.

The subsequent Illumina library preparation and sequencing yielded paired-end Hi-C reads of 150 bp at 75x-150x coverage. Scaffolding was carried out with contigs that were obtained using Canu and strain-specific Hi-C data. For scaffolding, the Hi-C reads were first mapped to each contig using BWA-MEM 0.7.17 (Li, 2013). Alignments were then processed with the Arima Genomics pipeline (Arima Genomics, 2019). Scaffolding was performed using SALSA 2.2 (Ghurye *et al*., 2017, 2019) and subsequent manual curation was guided by Hi-C contact maps. For strain C2, SALSA scaffolding resulted in 202 scaffolds from 325 contigs and 15 contigs were broken based on Hi-C contact map information. For strain A1, SALSA resulted in 107 scaffolds from 276 contigs and 14 contigs were broken based on Hi-C contact map information. For strain 4401, SALSA resulted in 108 scaffolds from 306 contigs and 13 contigs were broken based on Hi-C contact map information. For strain B3, SALSA resulted in 138 scaffolds from 314 contigs and 13 contigs were broken based on Hi-C contact map information. In the strain DAOM-197198 SALSA resulted in 153 scaffolds from 210 contigs, and one contig was broken based on Hi-C contact map information.

Gene annotation was achieved by using Funannotate (version 1.7.4) (https://zenodo.org/record/2604804#.YPm336iSnIU) on repeat-masked genome assemblies. Genome annotations were performed using RNA-seq reads and protein models for the strains DAOM197198, A1, B3 and C2 available from the Joint Genome Institute (JGI; (Chen *et al*., 2018b)). For the strain 4401, genome annotation was carried by combining EST, RNA-seq and models from the strains DAOM197198, C2 and B3. The quality of annotations was evaluated using the Benchmarking Universal Single-Copy Orthologs (version 4.0, database Obd_10) (Simão *et al*., 2015).

Conserved protein domains were predicted using Pfam v.27 (El-Gebali *et al*., 2019). SignalP 4.1 (-t euk -u 0.34 -U 0.34)(Petersen *et al*., 2011) and TMHMM 2.0(Krogh *et al*., 2001) were used to predict secreted proteins. A protein was called secreted if it was predicted to have a signal peptide and but no transmembrane domains. Effector candidates were predicted with EffectorP 3.0(Sperschneider & Dodds, 2021). De novo repeats were predicted with RepeatModeler 2.0.0 and the option -LTRStruct (Flynn *et al*., 2020). These were merged with the RepeatMasker repeat library and RepeatMasker 4.1.0 was run with this combined repeat database (Smit *et al*.). Transposable element locations were extracted from the Repeatmasker output file using an R script and simple repeats, unknown and low complexity repeats, satellites, tRNA, snRNA and rRNAs were filtered out from the output file. PfamScan (Hancock & Bishop, 2004; Li *et al*., 2015) was used with default parameters to identify the protein domain annotations in all strains. To create the Pfam heatmaps, the domain numbers were counted for each *R. irregularis* strain or A/B compartment, and a t-test conversion was made to highlight the protein domain abundances for each category.

We used mash dist (version 2.2.0) (Ondov *et al*., 2016) for *k-mer* distance calculations and dnadiff from the MUMmer (Kurtz *et al*., 2004; Marçais *et al*., 2018) suite for structural variation calculations between strains and compartments. Orthology analyses were made by FastOrtho (Wattam *et al*., 2014), using following parameters: 50% identity and 50% coverage using protein sequences of all five assemblies. All karyoplots were produced using KaryoploteR (Gel & Serra, 2017). All genome data and reads newly obtained are available in Genbank under the BioProject PRJNA748024.

### Identification of chromosome compartments and topologically associated domains

We called A/B compartments and TADs with hicexplorer 3.6 (Ramírez *et al*., 2018) using the commands hicPCA and hicFindTADs, respectively. Inter- and intra-chromosomal Hi-C contact maps were produced with HiC-Pro 2.11.1 (Servant *et al*., 2015) (MAPQ=10) and hicexplorer 3.6 (Ramírez *et al*., 2018; Wolff *et al*., 2018, 2020; Winter *et al*., 2018) and these Hi-C contact maps were manually inspected to assign chromosomal regions to A/B compartments. Specifically, the regions along the each chromosomes carrying the same PCA1 values – i.e. positive or negative – were assigned to the same compartment. Following this assignment, compartments were manually inspected to investigate their proximity with other chromosomes. Those in close physical proximity with the smallest chromosomes were assigned to compartment A, while those carrying the counterpart PCA1 signal were assigned to compartment B. This process was manually repeated for all 33 chromosomes individually to separate the chromosomal regions into separate A/B compartments. Bedtools (Quinlan & Hall, 2010) was used to assess overlap between genomic features such as genes/repeats and compartments.

### Methylation and gene expression analyses

Megalodon (version 2.2.9) was used for methylation calling using the high accuracy parameters (dna_r9.4.1_450bps_modbases_dam-dcm-cpg_hac.cfg) configuration file. After methylation calling was completed for all five assemblies, CG positions and their methylation frequencies were extracted from the 5mC bedfile output. For RNA-seq analyses, Salmon v.1.3.0 (Patro *et al*., 2017) was used to align clean RNA-seq reads to the transcripts and to estimate transcript abundances in each sample (salmon index –keepDuplicates and salmon quant – validateMappings). We used tximport and DESeq2 (Love *et al*., 2014) to assess gene differential expression (padj□<□0.1). The DAOM-197198 RNA-seq datasets used in this study were obtained from germinating spores (SRR1979300-SRR1979302), as well as intra-radical material isolated from *Medicago Trunculata* (SRR5644319-SRR5644324) and *Allium schoenopasum* (SRR5644331-SRR5644333).

## Results

### Assembly and annotation of Rhizophagus irregularis chromosomes

Basecalled and polished ONT data from four homokaryotic strains (A1, B3, C2 and 4401) were assembled using Canu (Koren *et al*., 2017), and then scaffolded using strain-specific Hi-C data. The same scaffolding procedures were performed on a previously published PacBio assembly of the model AMF strain DAOM197198 (Maeda *et al*., 2018). This approach resulted in an average of 137 scaffolds with an N50 score of 5 Mb, which were further curated into 33 chromosome-scale scaffolds in each strain, guided by Hi-C contact maps (**Table 1, Figure S1**).

**Table 1:** *R. irregularis* strains genome assembly and annotation statistics.

|  | **DAOM197198** | **A1** | **B3** | **C2** | **4401** |
| --- | --- | --- | --- | --- | --- |
| **Chromosome-scale scaffolds** |  |  |  |  |  |
| **Total size** | 147.2 Mb | 147.1 Mb | 146.8 Mb | 161.9 Mb | 146.8 Mb |
| **# of scaffolds** | 33 | 33 | 33 | 33 | 33 |
| **BUSCO** | 96% | 96% | 96% | 96.5% | 96.5% |
| **completeness** |  |  |  |  |  |
| **Repeat content** | 49.05% | 49.00% | 50% | 54.87% | 48.07% |
| **Number of genes** | 26,634 | 23,988 | 23,988 | 25,634 | 24,069 |
| **GC content** | 27.82% | 27.78% | 27.81% | 28.23% | 27.81% |
| **Unplaced contigs** |  |  |  |  |  |
| **Total size** | 2.6 Mb | 0.5 Mb | 1.3 Mb | 0.4 Mb | 0.9 Mb |
| **# of contigs** | 104 | 29 | 42 | 12 | 25 |
| **Assembly N/L50** | 29/31.4 Kb | 7/25.5 Kb | 12/36.5 Kb | 4/50.8 Kb | 7/45.7 Kb |

When combining the five strains, we found telomeres on both ends for 32 of the 33 chromosomes (only chromosome 2 has telomeres on only one end in all strains). Genome sizes are very similar in the model strain DAOM197198, A1, B3 and 4401 at 146-147 Mb, while the C2 strain harbors a larger genome at 162 Mb, consistent with published flow cytometry analyses (Ropars *et al*., 2016). For all five strains, the number of unplaced contigs is small (12-104 contigs at size 0.4-2.6 Mb) and all unplaced contigs are short indicating that the chromosome assemblies are near-complete (**Table 1, Figure S1**).

We identified an average of 76 families of repeats and known transposable elements (TE) encompassing on average 50.2% of the genome (**Figure S2**). When only TEs are considered, repeat density varies across chromosomes of different strains. For DAOM197198, A1 and B3 assemblies, chromosome 2 has the highest TE density (17.2%, 14.8% and 14.8% respectively), whereas the highest TE density is chromosome 33 for the C2 assembly (20.8%), and chromosome 23 for the 4401 assembly (15.6%). The chromosome with the lowest TE density is chromosome 20 for A1 and B3 (8.5% and 9.4%, respectively), chromosome 28 for DAOM197198 (9.2%), chromosome 3 for C2 (11.9%) and chromosome 8 for 4401 (7.3%).

Gene annotation identified between 23,693 to 26,820 genes, in line with past work based on fragmented datasets (Chen *et al*., 2018b), and BUSCO completeness averages 96% (**Table 1**). Gene density varies among chromosomes, with chromosome 32 always having the highest density while chromosome 15 the lowest (**Figure S1**). A hallmark of AMF is that they carry divergent rRNA operons within their genome. In DAOM197198 one rRNA operon is found on chromosome 9, two operons co-locate on chromosomes 23 and 28, and five are present in chromosome 18 (**Figure S1)**. The putative AMF mating-type locus (Ropars *et al*., 2016) is located on chromosome 11 in all strains (**Figure S1**).

### Within species chromosome diversity in R. irregularis

Consistent with previous reports on fragmented assemblies (Chen et al., 2018b), the *R. irregularis* gene content is divided into genes shared by all strains (core genes) and genes shared by only a few strains or strain-specific (dispensable) (**Figure S3a**), and on average only 55.8% of *R. irregularis* genes are core (Chen *et al*., 2018b). Our analyses confirm that within species variability affects the number of rRNA operons (**Figure 1a, Table S1**) (Corradi *et al*., 2007), and each strain carries a distinct abundance of protein domains (**Figure S3b**) (Chen *et al*., 2018b).

**Figure 1:**
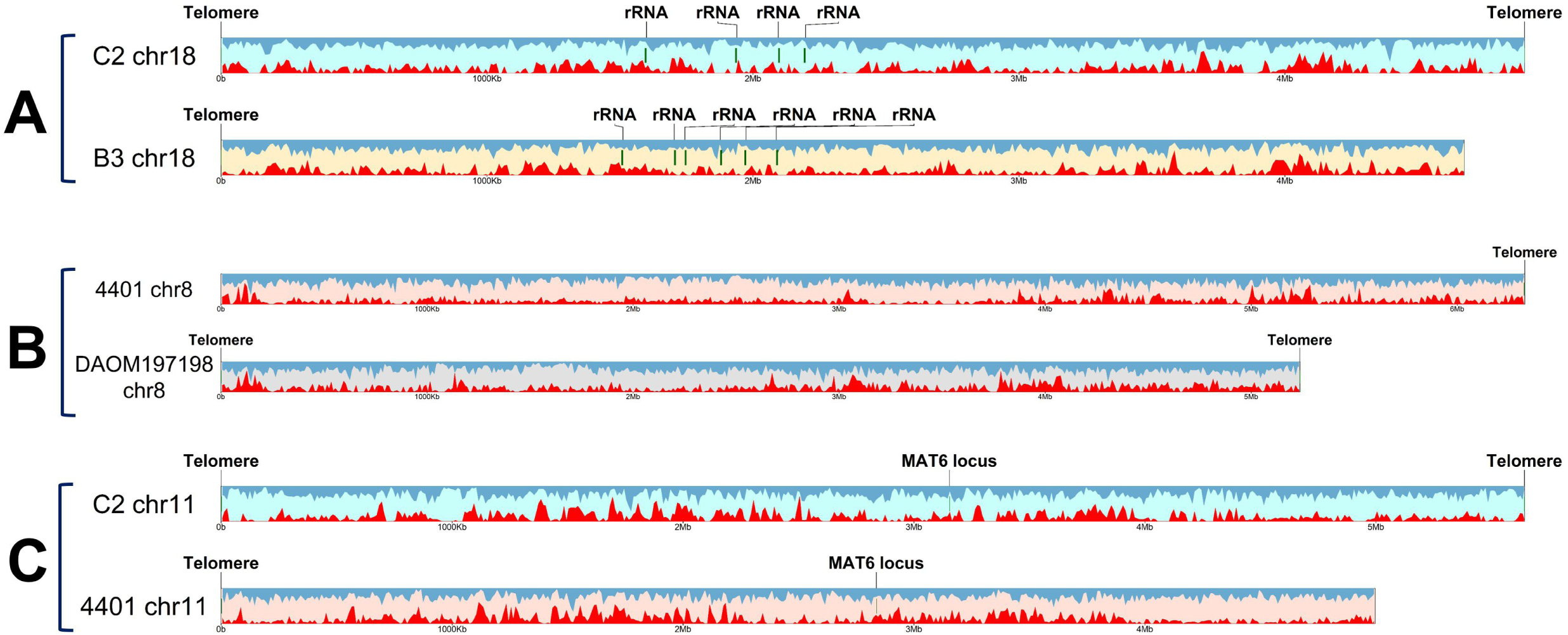
Examples of variability in the size, and gene content and density between the homologous chromosomes of R. irregularis strains analyzed in this study. Red density plot shows gene density, and blue density plot shows repeat density. The colors of karyoplots indicate different strains. **A) Homologous chromosomes contain different rRNA operon copy numbers**. Chromosome 18 of C2 contains only four operon copies, whereas strain B3 carries six copies on the same chromosome. **B) Homologous chromosomes vary greatly in size**. Size difference between chromosome 8 of 4401 strain and chromosome 8 of DAOM197198 strain is over 1.1 Mb. **C) Chromosomes carrying MAT locus also vary in size**. C2 and 4401 strains carry the same MAT type, MAT6. However, chromosome sizes still differ by 700 kb.

Homologous chromosomes often differ in size between strains. For example, the largest chromosome of the 4401 strain is chromosome 8 at 6.3 MB in size but this chromosome is only 5.2 MB in DAOM197198 (**Figure 1b**). The chromosome with the putative MAT-locus (Chromosome 11) varies in size from a maximum of 5.6 Mb in C2 to a minimum of 5 Mb in 4401, even though they carry the same mating type (**Figure 1c**). We used a *k*-mer distance measure to assess relatedness of strains and their chromosomes. Overall, the strain DAOM197198 is related to the strain 4401, strains B3 and A1 are related, and C2 is the most distant (**Figure S4**). However, this is not consistent across all chromosomes. For example, chromosome 8 shows higher levels of similarity between strain DAOM197198 and the strains B3 and A1 while chromosome 25 shows higher levels of similarity between strain 4401 and the strains B3 and A1 (**Figure S4**). These results support past reports of phylogenetic incongruence and inter-strain genetic exchange in *R. irregularis* (Riley *et al*., 2014; Chen *et al*., 2018b).

### The chromosomes of Rhizophagus irregularis form a two-compartment genome

Hi-C sequencing also generated direct, quantitative evidence of the 3D nuclear organization in five *R. irregularis* strains. For the strains DAOM197198, A1 and C2, this revealed that chromosomes form a distinct ‘checkerboard’ pattern that defines the long-range interactions that form two nuclear A/B compartments in eukaryotes, including fungi (Fortin & Hansen, 2015; Kim *et al*., 2017; Spielmann *et al*., 2018; Winter *et al*., 2018; Falk *et al*., 2019) (Figure 2, **Figures S5-10**).

**Figure 2:**
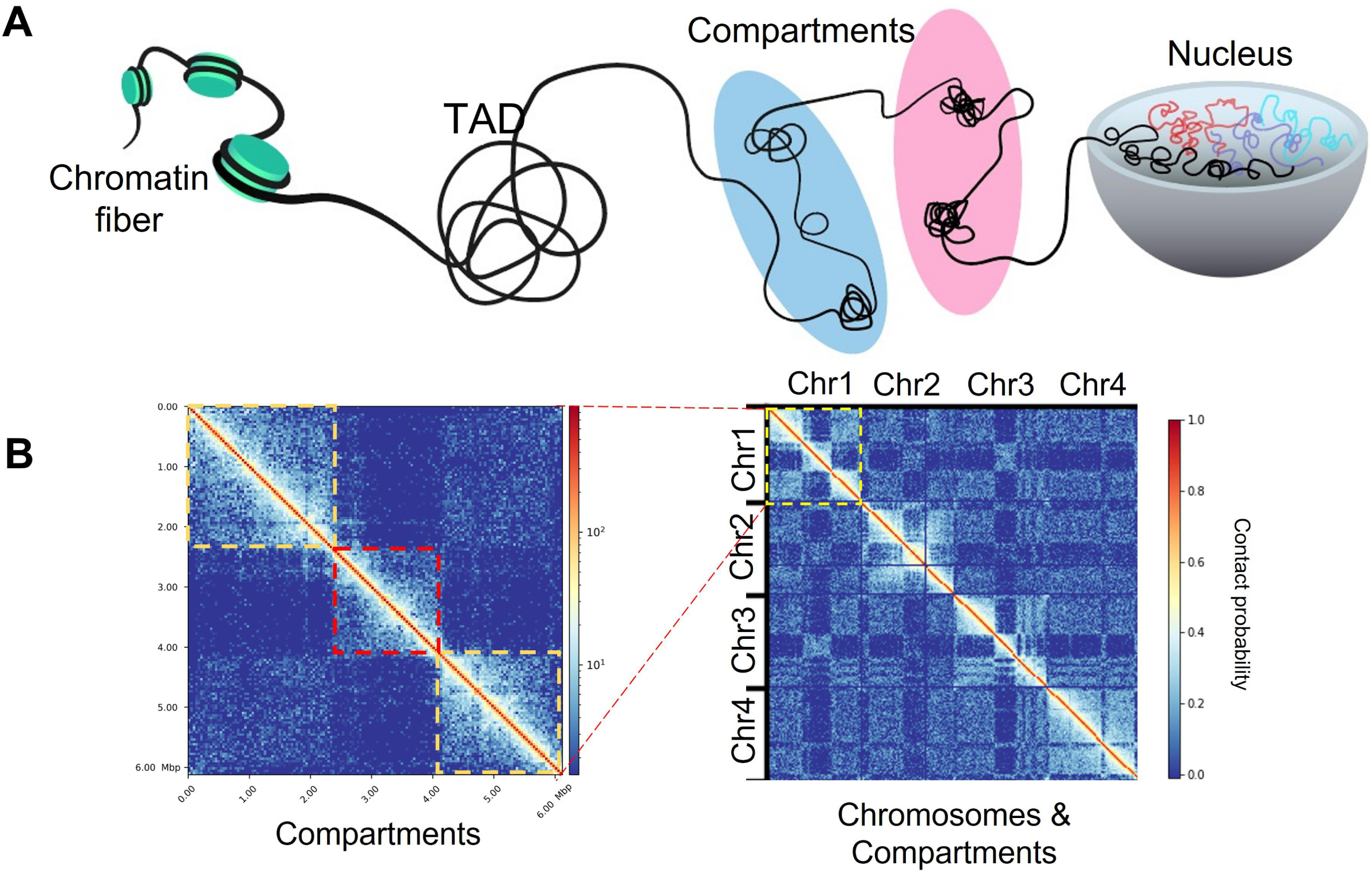
Chromatin folding inside of the *R. irregularis* nucleus and visualization of intra- and inter-chromosomal physical interactions in Hi-C maps. **A) Schematic showing the compaction of chromatin fibers inside of a nucleus**. Chromatin fibers physically interact with each other and may fold into regions called compartments and topologically associated domains. **B) Hi-C contact maps showing the compartmentalization in the chromosome 1 of the *R. irregularis***. In these heat maps, created to demonstrate contact frequencies throughout the genome, genome coordinates are represented on both axes. In an individual chromosome, regions that interact with each other result in an increase in contact frequencies and reveals the compartmentalized nature of the chromosome. These bright squares highlight increased contact frequency within and between chromosomes, are further analyzed to group them into euchromatin or heterochromatin compartments. Left: For chromosome l, the compartment shown in the red square belongs to compartment A, and is surrounded by two B-compartment regions shown in orange squares. Right: when the interactions of several chromosomes are analyzed, a “checkered” pattern indicates the genome arrangement of the euchromatin or heterochromatin compartments of *R. irregularis*. Telomeres are clearly visible in the Hi-C maps at the tip of each chromosome.

In the model strain DAOM197198, the A-compartment has a total size of 58.1 Mb while B-compartment is 73.5 Mb, and respectively each contains 96 and 83 TADs. Chromosome varies in the degree of compartmentalization – e.g. chromosome 2 has consistent swaps between A- and B-compartment, while the A-compartment dominates in the smallest chromosomes (chr30 – 33) **(Figure S6-10)**. Although *Rhizophagus irregularis* Hi-C contact maps did not show easily recognizable centromere hotspots found in rust fungi (Sperschneider *et al*., 2021), these reveal telomere-to-telomere interactions that define all chromosomes boundaries and confirm our assembly. Changes in Hi-C contacts between homologous chromosomes of different strains are also linked with the emergence of large strain-specific repeat and gene expansions (**Figure S11a**), or inversions (**Figure S11b**,**c**,**d**).

In the model strain DAOM197198, the A-compartment is more gene dense with 2.55 genes and 18.21 repeats per 10 Kb, while the B-compartment carries an average of 1.74 genes and 18.9 repeats per 10 Kb (**Figure 3a, Table S2)**. The A-compartment also codes for 90% of BUSCO proteins, and double the number of core genes (**Figure S12**), while B-compartment carries most dispensable genes and many highly expanded families – e.g. Tyrosine Kinases, the tetratricopeptide repeat Sel-1, the homodimerization BTB. The large high mobility box (HMG) gene family is, however, predominant in the B-compartment (**Figure 3b**). Remarkably, while the B-compartment is less gene-dense, it is enriched for predicted secreted proteins as well as for candidate apoplastic and cytoplasmic effectors and carries the cytoplasmic *R. irregularis* effector SP7 on chromosome 1 (Kloppholz *et al*., 2011).

**Figure 3:**
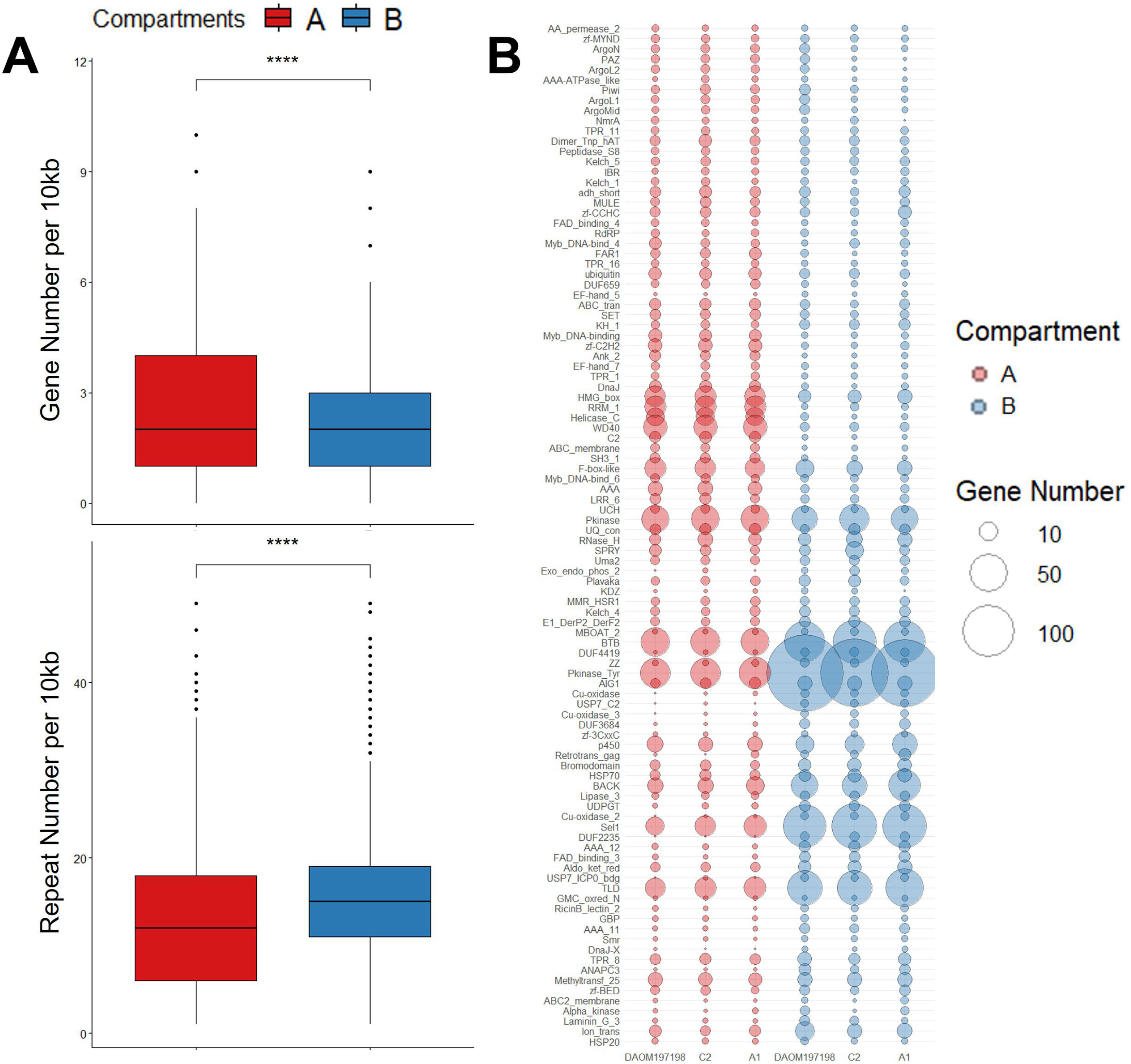
A/B compartments have different gene and repeat densities, and their genes contain different pfam domains. **A) The two compartments carry significantly different gene and repeat densities.** The boxplots show the number of gene and repeats in 10kb intervals. T-test shows that compartment A has significantly higher gene density, and significantly lower repeat density compared to compartment B. **B) Gene numbers that carry specific Pfam domains also vary between compartments**. Circles highlight the total of number of genes carrying specific Pfam domains in located in compartment A (red circles) or B (blue circles), with the size of the circle being proportionate to the number of genes that carry that domain. Inter-strain variability in pfam domains is also evident within each compartment.

We hypothesized that the distinct gene and repeat densities result in different structural rearrangement rates between A/B compartments. We measured the degree of structural variation and sequence identity of each compartment, and found that the sequence alignments between strains in the B-compartment are shorter and have lower sequence identity than sequence alignments in the A-compartment. The B-compartment also contains more structural variation (**Table S3, Figure S13**).

### Gene expression and methylation in chromosomal compartments

In other organisms, euchromatin (A compartments) is generally transcriptionally active while heterochromatin (B compartments) are repressed (Szabo *et al*., 2019; Zheng & Xie, 2019; Jerković *et al*., 2020). To test if this dichotomy also applies to AMF, we investigated gene expression between compartments in DAOM197108 using three high quality RNA-seq datasets obtained from intra-radical hyphae in symbiosis with *Allium*, arbuscocytes in symbiosis with *Medicago* and germinating spores (Kamel *et al*., 2017; Zeng *et al*., 2018).

In *R. irregularis*, genes in the A compartment have, on average, significantly higher expression levels than those in the B compartment (**Figure 4a**). However, this compartment also has lower expression levels in the *in planta* conditions (*Allium*: 57.3 TPMs; *Medicago*: 56 TPMs) than in germinated spores (62.4 TPMs). Remarkably, the inverse pattern is observed for genes in the B-compartments, which have higher expression levels in the *in planta* conditions (*Allium*: 16.8 TPMs; *Medicago*: 19.5 TPMs) than in germinated spores (11.2 TPMs), and genes encoding secreted proteins are significantly up-regulated in both *in planta* conditions in the B-compartment, but not in the A-compartment (**Figure 4b**). We also aimed to investigate the expression of these regions within TAD, and found that genes tend to be co-regulated when they are located in the same TAD in the A-compartment, while co-regulation within the same TAD was not observed for the B-compartment **(Figure S14)**.

**Figure 4:**
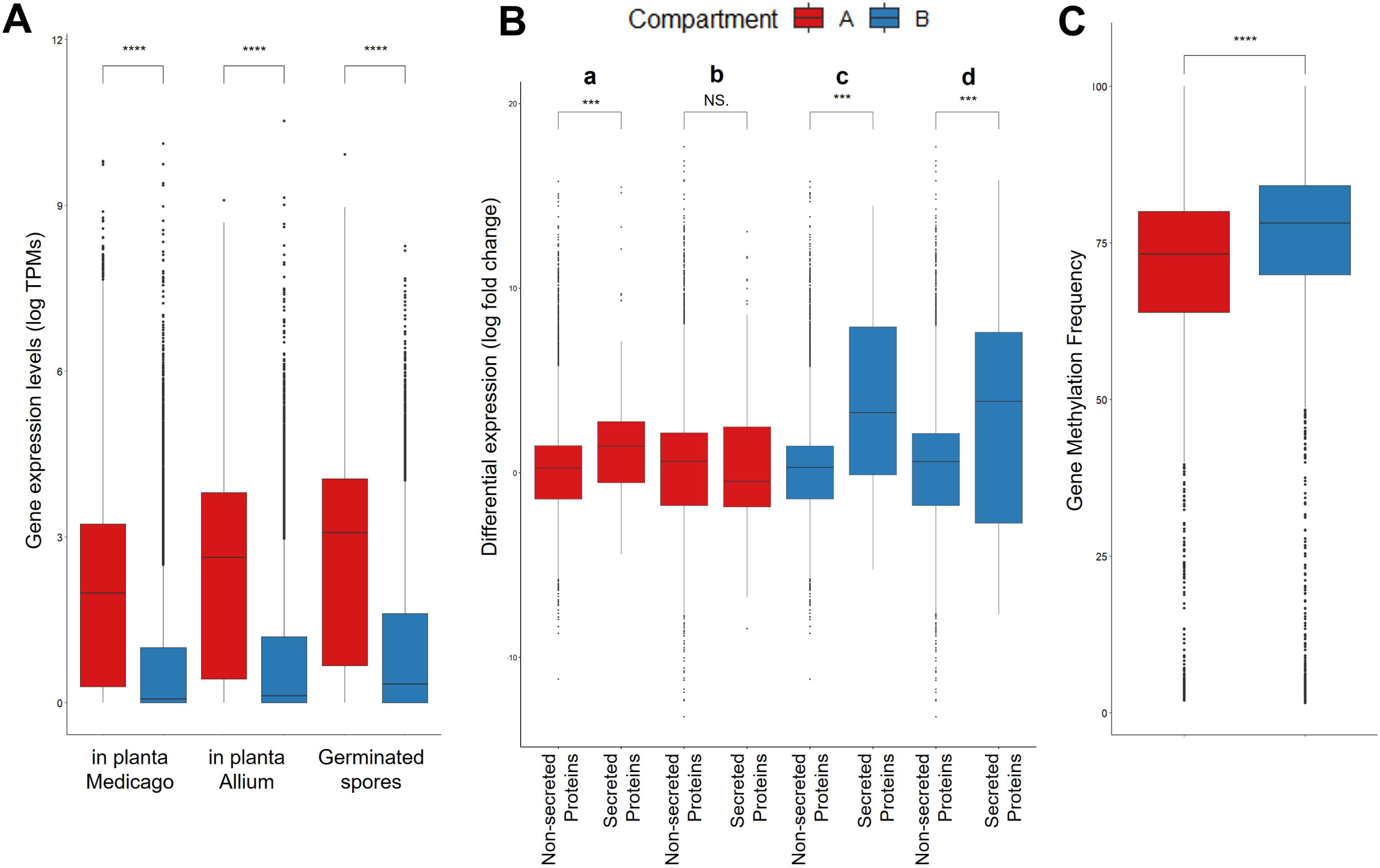
Gene expression and methylation analyses for the A/B compartments. T-test was conducted to show significance. **A) Genes in the A compartment show significantly higher expression levels measured in Transcripts Per Million (TPM) than genes in the B-compartment in all three conditions. B) Genes encoding secreted proteins are up-regulated in both in planta conditions in the B-compartment, but not in the A-compartment. a)** Compartment A shows significant upregulation of secreted proteins *in planta* Allium condition when compared to germinated spores. **b)** Compartment A does not show a significant secreted protein expression change when *in planta* Medicago expressions are compared to the expression of germinated spores. **c)** Compartment B shows significant upregulation of secreted proteins *in planta* Allium condition when compared to germinated spores. **d)** Compartment B shows significant upregulation of secreted proteins *in planta* Medicago condition when compared to germinated spores. **C) Gene median methylation frequencies of all methylated genes (methylation frequency median > 0) that are located in A/B compartments**. A compartment display significantly lower gene median methylation frequencies than the B compartment.

We used ONT sequencing to call methylated sites and test if the difference in the A/B compartment gene expression resulted from different methylation patterns (Lea *et al*., 2018; Dallaire, 2021). In DAOM197198 strain, CG dinucleotides are highly enriched for 5mC methylation – i.e. 11.88% for CG compared to 0.72 to 3.95% for other dinucleotides contexts (CA, CC, CT) – 6mA methylation is very low regardless of dinucleotide contexts (AA, AC, AG, AT: 0.01% to 0.06). This indicates that 5mC methylation in the CG dinucleotide context is the primary DNA methylation process active in this species.

Overall, the CpG sites are significantly more methylated in the A-compartment, particularly for highly methylated sites (methylation frequency > 80%) (**Figure S13a)**. Within the B compartment, 22.5% of all protein encoding genes are highly methylated compared to 18.4% in the A-compartment (**Figure 4c**). When TEs are considered, this is inversed with 67% of methylated TEs families (24 out of 35) being highly methylated in the A-compartment compared to only 25% for the B -compartment (9 out of 36) (**Figure S13b**). Among TEs that are compartment specific - e.g Ginger DNA 2, DNA Zisupton and LINE Penelope are only found in compartment B and the LINE -Tad, and LINE R1-LOA carried by the A-compartment – only those located in the A-compartment are methylated.

## Discussion

### A chromosome-level view of a model AMF pangenome

The present study confirms that the combined number of genes within the model AMF *R. irregularis* far exceeds that found in individual strains (Chen *et al*., 2018b; Mathieu *et al*., 2018b; Reinhardt *et al*., 2021). It also revealed that strains in this model species carry 33 homologous chromosomes (possibly the largest number identified in a fungus) and within-species variability in chromosome size and epigenetics (folding and methylation).

Size difference among homologous chromosomes affects all strains, regardless of their phylogenetic clustering (Savary *et al*., 2018; Chen *et al*., 2018b; Kokkoris *et al*., 2021) or MAT-locus identity – e.g. strains C2 and 4401 carry the MAT-locus-6 (as defined by (Ropars *et al*., 2016)) on chromosome 11 but this chromosome is 640 Kb larger in C2 compared to 4401. Despite this variability, all strains analysed carry the same number of homologous chromosomes, and future studies may reveal if this characteristic defines species boundaries in this taxonomically challenging taxon (Bruns *et al*., 2018).

### Chromosome compartments dictate AMF genome biology and evolution

Hi-C data revealed that AMF chromosomes separate into locations with euchromatin (transcriptionally active A-compartment) and heterochromatin (transcriptionally less active B-compartments) that regulate both the expression and evolution of distinct molecular functions. In the A-compartment, higher gene density and repeat methylation indicates a tighter control of repeat expansions that could be deleterious for a region that contain most core genes. In contrast, the B-compartment experiences higher gene and lower repeat methylation rates, which explains its reduced gene expression levels and higher rearrangement rates.

It was proposed that AMF and notorious plant pathogens (e.g. *Fusarium oxysporum, Verticillium dahlia*; *Zymoseptoria tritici*) evolved similar genomic strategies to cope with their evolving plant hosts and competitors (Reinhardt *et al*., 2021). The A/B compartments are reminiscent of the “two-speed genome architecture” reported in some fungal plant pathogens, where highly repetitive and rapidly rearranging genomic regions that express secreted proteins (i.e. analogous to B-compartments) separate from gene-rich locations that carry most core genes (i.e. analogous to A-compartments) (Mathieu *et al*., 2018a). Another similarity with plant pathogens is the presence in *R. irregularis* strains of many dispensable and lineage-specific genes. Despite this genomic resemblance, however, our analyses did not reveal the presence in *R. irregularis* of dispensable chromosomes that drive the adaptation of fungal plant pathogens to changing environments (Garmaroodi & Taga, 2007; Vlaardingerbroek *et al*., 2016; Bertazzoni *et al*., 2018).

### A role of a two-compartment genome in plant colonization?

We showed that the A/B compartments have distinct epigenetic signatures, and in line with mammalian studies, the A-compartment in *R. irregularis* is transcriptionally active whereas the B-compartment is repressed. However, we also found that this does not hold true for genes encoding secreted proteins and candidate effectors. During two *in planta* conditions, we observed increased up-regulation of genes encoding secreted proteins in the B-compartment, but not in the A-compartment. This suggests that the repressed state of the B-compartment might be relaxed during plant colonization, possibly induced through signals from the plants (Plett & Martin, 2012). In the fungal pathogen *Leptosphaeria maculans* epigenetic control mechanisms lead to effector gene regulation (Soyer *et al*., 2014). As our Hi-C data derives from extraradical mycelium, acquiring similar data *in planta* may show that intra-radical mycelium carries distinct chromosome conformations that lift repression for the *R. irregularis* B-compartment (Frantzeskakis *et al*., 2019; Torres *et al*., 2020).

## Conclusions

Combining long-reads with Hi-C sequencing demonstrates that model arbuscular mycorrhizal symbionts all carry 33 chromosomes with contents and sizes that varies significantly among strains. This supports the notion that AMF have pangenomes and, more generally, that conspecific strains should never be assumed to have identical genomes and genes (Malar C *et al*., 2021b).

Our work also uncovered a higher-order genome organization that governs AMF genome biology and evolution. Dispensable genes and the most expressed and regulated secreted proteins - i.e. those potentially involved in the molecular dialogue with the plant hosts - locate primarily in the rapidly evolving B-compartments, while highly expressed core genes and tightly regulated TADs are mostly restricted within the slowly evolving A-compartment. In other organisms, the A/B compartments change depending on the cell life-stage (Falk et al., 2019). Within this context, our findings raise the intriguing possibility that similar changes also occur depending on the fungal and plant symbiotic status, and open avenues to identify the epigenetic mechanisms that generate and modify chromosome compartments during host-microbe interactions.

It will be now interesting to examine how this work extrapolates to AMF dikaryons, in particular how these strains compartmentalize their co-existing parental genomes and if these also significantly differ in gene content and size. Phasing parental genomes with high-quality Hi-C data is a requirement to fully address these questions, and determine how co-existing genomes regulate nuclear dynamics in these strains (Serghi *et al*., 2021; Kokkoris *et al*., 2021).

## Acknowledgments

We thank Benjamin Schwessinger and Daniel Croll for their helpful comments on an earlier version of this manuscript, and Christophe Roux for providing information about publicly available, high-quality RNA-seq data. Our research is funded by the Discovery program of the Natural Sciences and Engineering Research Council (RGPIN2020-05643), a Discovery Accelerator Supplements Program (RGPAS-2020-00033). N.C. is a University of Ottawa Research Chair in Microbial Genomics. J.S. is supported by an Australian Research Council (ARC) Discovery Early Career Researcher Award (DE190100066), and E.C. and W.I. were supported by JSPS Postdoctoral Fellowships for Research in Japan and JSPS KAKENHI Grant Number 19F19089.

## Supplemental figures and tables legends

**Figure S1: *R. irregularis* C2, A1, B3 and 4401 strain chromosomes**. The length of all 33 chromosomes is indicated in the karyoplots. Telomere repeats, MAT locus and small subunit rRNA locations are marked. Gene densities are shown in red, and repeat densities are shown in blue color.

**Figure S2:** Bubble plot showing 10 most abundant transposable elements and their distribution in five assemblies of *R. irregularis* strains.

**Figure S3: Gene orthologies and Pfam domain numbers in five strains of *R. irregularis*. a)** Protein orthologs of A1, B3, C2, DAOM-197198 and 4401 strains. **b) Pfam protein domain abundance comparison between five *R. irregularis* strains**. After the 100 most abundant Pfam protein domains were selected, the frequency values were transformed using a t-test. This way, relative enrichment (red) and relative depletion (blue) were shown.

**Figure S4: Hierarchical clustering of *k*-mer distance estimations between chromosomes of the strains**.

**Figure S5: Heat map showing Hi-C contact frequencies between 33 chromosomes for strain DAOM-197198, C2, A1, B3 and 4401**. Regions having higher contact frequencies are found in close contact in the nucleus, whereas regions having fewer contact frequencies are far away from each other.

**Figure S6: Heat map showing Hi-C contact probabilities of individual chromosomes of strain DAOM-197198**. Chromosomal regions having higher contact probabilities are found in close contact with each other, whereas regions having fewer contact probabilities are far away from each other. PCA eigenvectors were calculated for 10kb intervals and were plotted under the chromosome contact maps.

**Figure S7: Heat map showing Hi-C contact probabilities of individual chromosomes of strain C2**. Chromosomal regions having higher contact probabilities are found in close contact with each other, whereas regions having fewer contact probabilities are far away from each other. PCA eigenvectors were calculated for 10kb intervals and were plotted under the chromosome contact maps.

**Figure S8: Heat map showing Hi-C contact probabilities of individual chromosomes of strain A1** Chromosomal regions having higher contact probabilities are found in close contact with each other, whereas regions having fewer contact probabilities are far away from each other. PCA eigenvectors were calculated for 10kb intervals and were plotted under the chromosome contact maps.

**Figure S9: Heat map showing Hi-C contact probabilities of individual chromosomes of strain B3** Chromosomal regions having higher contact probabilities are found in close contact with each other, whereas regions having fewer contact probabilities are far away from each other. PCA eigenvectors were calculated for 10kb intervals and were plotted under the chromosome contact maps.

**Figure S10: Heat map showing Hi-C contact probabilities of individual chromosomes of strain 4401** Chromosomal regions having higher contact probabilities are found in close contact with each other, whereas regions having fewer contact probabilities are far away from each other. PCA eigenvectors were calculated for 10kb intervals and were plotted under the chromosome contact maps.

**Figure S11: Strain-specific chromosomal differentiation events can be observed in Hi-C contact maps**. Differentiation events that can be observed on contact maps and synteny plots were shown with an orange arrow. Repeat densities of the chromosomes are shown in blue color, and the synteny blocks are shown in purple. **a) Strain-specific repeat expansion found in chromosome 8 of strain 4401**. A) A repeat expansion event can be observed in chromosome 8 of 4401 strain, between regions 1Mb and 3Mb. As the same repeats get expanded, the contact of Hi-C reads in that region increase, leading to the increased brightness. B) Synteny blocks between chromosome 8 from DAOM-197198 and 4401 shows absence of synteny blocks in the repeat expansion region. In the regions showing no synteny between the two strains, DAOM-197198 strain has 157 genes and 1240 repeats whereas 4401 strain has 389 genes and 2799 repeats. **b) Strain specific inversion event in chromosome 18 of strain C2**. A) An inversion event can be observed in chromosome 18 of C2 strain, around 2Mb region. This differentiation event also causes an abnormality in the contact map around that region. B) Synteny blocks between chromosome 18 from DAOM-197198 and C2 show multiple inversion events. **c) Strain specific inversion event in chromosome 17 of strain A1**. A) An inversion event can be observed in chromosome 17 of A1 strain, before 1 Mb region. This differentiation event also causes an abnormality in the contact map around that region. B) Synteny blocks between chromosome 17 from DAOM-197198 and A1 show multiple inversion events. **d) Strain specific inversion event in chromosome 18 of strain B3**. A) An inversion event can be observed in chromosome 18 of B3 strain, around 2Mb region. This differentiation event also causes an abnormality in the contact map around that region. B) Synteny blocks between chromosome 18 from DAOM-197198 and B3 show multiple inversion events.

**Figure S12: Distribution of core and dispensable genes in compartments A and B**. Protein orthologs of A1, C2 and DAOM-197198 strains were identified with FastOrtho (Wattam *et al*., 2014), using following parameters: 50% identity and 50% coverage. The genes were then grouped into A and B compartments. Core and core duplicated gene numbers were added up in each compartment to represent the “core group” count, and dispensable, specific and specific duplicated gene numbers were added up to make up the “dispensable group” count.

**Figure S13: A)** For each A/B compartment, the median methylation frequencies are collected and their distribution is plotted. The A compartment display higher median methylation frequencies than the B compartment. Highly methylated CpG sites (median methylation frequency > 80%) occur more frequently in the A compartment. **B)** CpG methylation frequencies of transposable elements in compartments A and B. Individual CpG methylation frequencies of transposable elements found in compartments A and B are shown as green dots. Methylation medians are shown as red dots.

**Figure S14: Example of inversions in synteny blocks between chromosome 13 of DAOM-197198, C2 and A1 strains**. The colors on the karyoplots represent the compartments. Compartment A is represented by blue color, whereas compartment B is represented by red color. Synteny blocks are created by MCScanx(Wang *et al*., 2012), and inversions are highlighted. Orange links show inverted synteny blocks between DAOM-197198 and A1, and green links show inverted synteny blocks between DAOM-197198 and C2. Majority of the inversions are observed in compartment B.

**Table S1:** Location and number of ribosomal RNA operons within and across strains analyzed in this study.

**Table S2: A/B compartment properties in the model strain DAOM-197198**. The A-compartment is more gene-dense, while the B-compartment is more repeat-rich. However, the B-compartment has significantly more secreted proteins, candidate apoplastic effectors and candidate cytoplasmic effectors than the A-compartment (Fisher’s exact test: * indicates p < 0.05).

**Table S3: Comparison of A and B compartment similarity in DAOM-197198, C2 and A1 strains**. Overall, A compartment is more conserved with higher similarity between strains compared to B compartment.

